# A rare opportunist, *Morganella morganii*, decreases severity of polymicrobial catheter-associated urinary tract infection

**DOI:** 10.1101/754135

**Authors:** Brian S. Learman, Aimee L. Brauer, Kathryn A. Eaton, Chelsie E. Armbruster

**Author notes:** Corresponding author: Chelsie Elizabeth Armbruster, 955 Main Street, Room 5218, Buffalo, NY 14203.

## Abstract

Catheter-associated urinary tract infections (CAUTIs) are common hospital-acquired infections and frequently polymicrobial, which complicates effective treatment. However, few studies experimentally address the consequences of polymicrobial interactions within the urinary tract, and the clinical significance of polymicrobial bacteriuria is not fully understood. *Proteus mirabilis* is one of the most common causes of monomicrobial and polymicrobial CAUTI, and frequently co-colonizes with *Enterococcus faecalis, Escherichia coli, Providencia stuartii*, and *Morganella morganii. P. mirabilis* infections are particularly challenging due to its potent urease enzyme, which facilitates formations of struvite crystals, catheter encrustation, blockage, and formation of urinary stones. We previously determined that interactions between *P. mirabilis* and other uropathogens can enhance *P. mirabilis* urease activity, resulting in greater disease severity during experimental polymicrobial infection. Our present work reveals that *M. morganii* acts on *P. mirabilis* in a contact-independent manner to decrease urease activity. Furthermore, *M. morganii* actively prevents urease enhancement by *E. faecalis, P. stuartii*, and *E. coli.* Importantly, these interactions translate to modulation of disease severity during experimental CAUTI, predominantly through a urease-dependent mechanism. Thus, products secreted by multiple bacterial species in the milieu of the catheterized urinary tract can directly impact prognosis.

## Introduction

Urinary catheters are common in health care settings, estimated to be utilized in ∼60% of critically ill patients, 20% of surgical unit patients, and 10% of nursing homes residents (1–3). The vast majority of individuals with an indwelling urinary catheter will experience bacteriuria and may progress to catheter-associated urinary tract infection (CAUTI) (1). CAUTI is the most common healthcare-associated condition in the United States, and carries an estimated economic burden approaching $1.7 billion annually (4).

Bacteriuria in catheterized individuals rapidly becomes polymicrobial, involving combinations of *Proteus mirabilis, Enterococcus faecalis*, *Escherichia coli, Pseudomonas aeruginosa, Klebsiella pneumoniae, Providencia stuartii, Morganella morganii, Citrobacter* species, *Staphylococcus* species, and *Streptococcus* species (1, 2, 5). It is estimated that 31-86% of CAUTIs are also polymicrobial (6–14); however, urine specimens are often suspected of harboring periurethral or vaginal microbiota if multiple colony types are observed, particularly if Gram-positive organisms are present, and are therefore often dismissed as “contamination” (15, 16). Thus, the clinical significance of polymicrobial bacteriuria is not fully understood.

There are numerous experimental and clinical examples of polymicrobial colonization causing more severe disease than infection with a single bacterial species (monomicrobial), including periodontitis, abdominal abscesses, actinomycosis, chronic wounds, otitis media, pneumonia, cystic fibrosis, inflammatory bowel disease, uncomplicated urinary tract infection, and catheter-associated urinary tract infection (17–20). In many cases, polymicrobial interactions promote an increased level of colonization for one or both of the bacterial species. It is also possible for a species that is generally perceived as a commensal organism to become an “accessory pathogen” by supporting or enhancing the virulence of a more traditional pathogen (21). In contrast to these disease-promoting interactions, other polymicrobial interactions attenuate disease severity or exclude colonization by traditional pathogens (21). Considering the prevalence of polymicrobial bacteriuria and CAUTI in catheterized individuals, exploration of polymicrobial interactions and their impact on disease severity is necessary.

Our recent analysis of 182 clinical CAUTIs from a 3-year study at 12 nursing homes identified *P. mirabilis* as the most common cause of CAUTI (26% of cases), followed by *Enterococcus* species (21%) and *Escherichia coli* (20%) (22). Thirty-one percent of the 182 CAUTIs were polymicrobial, and the most common bacterial combination was *P. mirabilis* with *E. faecalis*, which is in agreement with prior work regarding the composition of polymicrobial CAUTI (1, 5, 12–14, 22, 23). Notably, *E. faecalis, M. morganii*, and *P. stuartii* were significantly more prevalent during polymicrobial CAUTI than monomicrobial infection, and each of these bacterial species has been reported to co-colonize catheters with *P. mirabilis* (1, 5, 12–14, 22, 24). These five species therefore represent important constituents of the polymicrobial CAUTI environment.

*P. mirabilis* poses a significant challenge for effective CAUTI treatment as *Proteus* isolates are intrinsically drug resistant, exhibiting high tolerance to tetracycline and polymyxin, and clinical isolates are often resistant to aminoglycosides and fluoroquinolones (25). Furthermore, there are increasing reports of isolates producing extended-spectrum β-lactamases (ESBL) and carbapenemases (25–29), which threatens the utility of last-resort antibiotics and increases the mortality rate for *P. mirabilis* infection (30–32). *P. mirabilis* also acts as a “hub” species in catheterized nursing home residents, promoting colonization by additional multidrug resistance organisms (33) and providing protection from antibiotic treatment (34). Furthermore, *P. mirabilis* produces a potent urease enzyme that hydrolyzes the urea in urine to carbon dioxide and ammonia, thereby increasing urine pH and facilitating the precipitation of polyvalent ions and resulting in struvite crystals, catheter encrustation, blockage, and formation of urinary stones (urolithiasis) (35–37). Urolithiasis in humans and animal models of infection can elicit bladder obstruction and renal damage (35, 38, 39), which facilitate sepsis and bacteremia. Indeed, *P. mirabilis* is the causative agent in 13-21% of bacteremias experienced by nursing home residents, the majority of which are secondary to CAUTI (9, 40–45). However, there are examples of catheterized patients with prolonged colonization by *P. mirabilis* who do not experience catheter blockage or urolithiasis (12, 46), indicating that the magnitude of struvite crystal formation is likely affected by other factors within the urinary tract environment.

In our prior investigations of *P. mirabilis* urolithiasis and pathogenicity, we determined that numerous common urinary tract colonizers are capable of enhancing *P. mirabilis* urease activity during co-culture in urine *in vitro* (47). *E. faecalis, E. coli*, and *P. stuartii* were the most potent enhancers of *P. mirabilis* urease activity, and experimental coinfection of *P. mirabilis* with *P. stuartii* in a murine model of complicated UTI resulted in higher urinary pH, a greater incidence of urinary stones, and increased disease severity, all of which were dependent on the presence of a functional urease operon in *P. mirabilis* (47, 48). Thus, the concomitant presence of certain urease-enhancing organisms may correspond to likelihood of developing struvite crystals and urolithiasis during *P. mirabilis* CAUTI. In contrast, *M. morganii* was the only species tested that did not enhance *P. mirabilis* urease *in vitro*, and may have even dampened activity (47), suggesting that this species may be able to decrease risk of struvite formation during polymicrobial colonization and therefore possibly reduce disease severity.

Like *P. mirabilis, M. morganii* is a urease-positive organism. However, it produces a urease enzyme that is distinct from that of *P. mirabilis* (49), and the urine pH change mediated by *M. morganii* urease activity rarely results in development of struvite crystals or catheter blockage (24, 50–52). Interestingly, *M. morganii* is more commonly isolated from unobstructed catheters than blocked catheters in patients catheterized long-term (12), and pre-colonization of a catheter by *M. morganii* can actually reduce blockage and encrustation by *P. mirabilis in vitro* (24), albeit transiently. Taken together, these data support the hypothesis that *M. morganii* may have the capacity to antagonize *P. mirabilis* urease activity and reduce the incidence of struvite crystal formation and urolithiasis during co-culture.

In this study, we tested the urease-modulatory capability of *M. morganii* and its impact on disease severity during dual-species and triple-species combinations. We show that *M. morganii* secretes products that decrease *P. mirabilis* urease activity in a contact-independent manner, and that the urease-dampening capability of these secreted products can prevent urease enhancement by *E. faecalis, P. stuartii*, and *E. coli.* Using a murine model of polymicrobial CAUTI, we further demonstrate that coinfection of *P. mirabilis* with *E. faecalis* dramatically increases disease severity, predominantly due to enhancement of *P. mirabilis* urease activity, and that the addition of *M. morganii* to this polymicrobial combination mitigates disease severity. *M. morganii* was also capable of mitigating cytotoxicity *in vitro.* Thus, identification of the *M. morganii* secreted products that reduce *P. mirabilis* pathogenicity may uncover a novel therapeutic strategy to prevent catheter encrustation, urolithiasis, tissue damage, and secondary bacteremia resulting from *P. mirabilis* catheter colonization.

## Results

### Uropathogens exhibit comparable competitive fitness during co-culture in dilute human urine

To determine if *P. mirabilis* exhibits cooperative or competitive behavior with *E. faecalis, E. coli*, or *M. morganii*, each bacterial species was cultured independently or co-cultured in dilute human urine and bacterial viability was assessed by differential plating at hourly timepoints until the cultures achieved stationary phase (Figure 1). The viability of *P. mirabilis* was not significantly impacted by co-culture with any of the other species (Figure 1A), and similar results were observed for *E. faecalis* (Figure 1B) and *M. morganii* (Figure 1C), indicating that these species do not exhibit competitive behavior during co-culture *in vitro*. In contrast, *E. coli* viability was unaffected by *M. morganii* but dramatically reduced during co-culture with *P. mirabilis* (Figure 1D). The reduction in viability was pH-dependent, as *E. coli* viability did not decrease during co-culture with a *P. mirabilis* urease mutant (Δ*ure*). Thus, *E. coli* is sensitive to the increase in urine pH that occurs due to *P. mirabilis* urease activity, but these species do not appear to exhibit any other competition *in vitro*.

**Figure 1.**
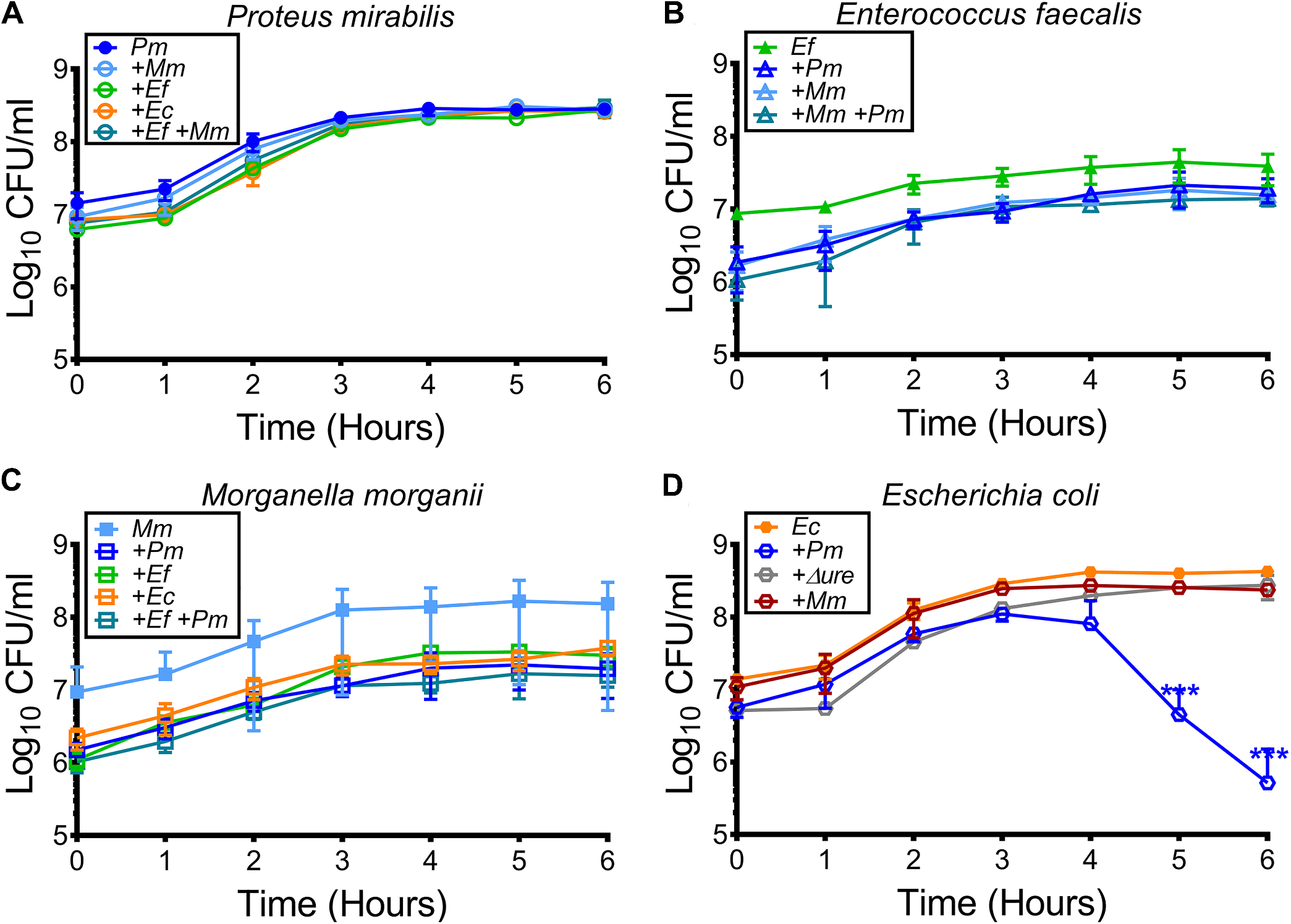
Growth of urinary tract isolates in human urine. *P. mirabilis* HI4320 (*Pm*), *E. faecalis* 3143 (*Ef*), *M. morganii* TA43 (*Mm*), and *E. coli* CFT073 (*Ec*) were cultured in rich laboratory media overnight, diluted 1:100 into filter-sterilized human urine, and incubated at 37°C with aeration for 6 hours for hourly assessment of growth and viability by determination of CFUs. (A) CFUs of *P. mirabilis* from monomicrobial and polymicrobial cultures, (B) CFUs of *E. faecalis* from monomicrobial and polymicrobial cultures, (C) CFUs of *M. morganii* from monomicrobial and polymicrobial cultures, and (D) CFUs of *E. coli* from monomicrobial and polymicrobial cultures, including during incubation with *P. mirabilis* Δ*ure*. Error bars represent mean ± SD for three independent experiments. Data were log_10_-transformed for statistical analyses, and significance was assessed by 2-way ANOVA with Dunnett’s multiple comparisons test relative to monomicrobial cultures. \*\*\**p*<0.001.

### M. morganii prevents enhancement of P. mirabilis urease activity by other bacterial species in a contact-independent manner

We previously demonstrated that numerous uropathogens, including *E. faecalis* (*Ef*)*, P. stuartii* (*Ps*), and *E. coli* (*Ec*), can enhance *P. mirabilis* (*Pm*) urease activity during co-culture in urine while *M. morganii* (*Mm*) appears to dampen activity (47). We therefore sought to determine if urease modulation by these species requires direct cell-cell contact or if *P. mirabilis* may be responding to factors present in cell-free supernatants. An example of the experimental approach is shown in Figure 2A. Briefly, each bacterial species of interest was incubated in filter-sterilized human urine for 90 minutes, centrifuged to pellet, and filter-sterilized. Cell-free supernatants were also generated from co-cultures of *M. morganii* with each of the other bacterial species for comparison to 50:50 mixtures of the monomicrobial culture supernatants. In addition, cell-free supernatant was generated from an isogenic urease mutant of *P. mirabilis* HI4320 (Δ*ure*) for use as a control for the potential impact of nutrient and cofactor depletion on total urease activity. Wild-type *P. mirabilis* HI4320 was then incubated in each of the cell-free supernatants, and urease activity was measured over a 120-minute time course via a colorimetric assay (Figure 2B and C). Urease activity was also quantified as the mean change in optical density per minute for the linear portion of the activity curve (Figure 2D).

**Figure 2.**
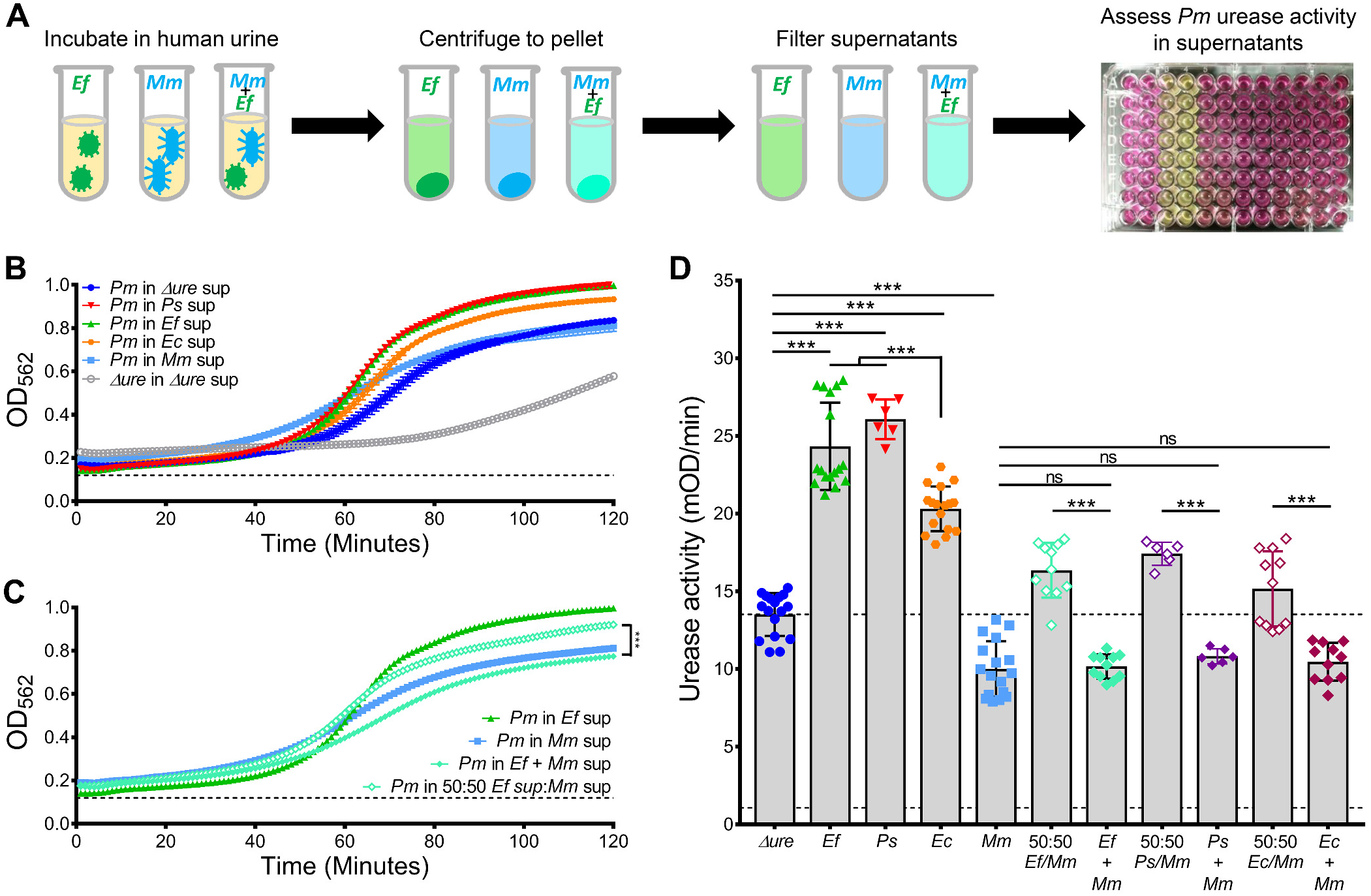
Urease activity of *P. mirabilis* during incubation in cell-free urine supernatants. An example of the experimental approach is displayed in panel A. Cell-free urine supernatants were generated from monomicrobial cultures of strains of interest (*P. mirabilis* Δ*ure, E. faecalis* 3143, *E. coli* CFT073, and *M. morganii* TA43) by 90-minute incubation in filter-sterilized human urine. Supernatants were also generated from co-cultures of *M. morganii* with each of the other bacterial species. *P. mirabilis* HI4320 was then incubated in each of the cell-free urine supernatants supplemented with excess urea, and urease activity was monitored at 60-second intervals over 120 minutes. (B) *P. mirabilis* urease activity in each of the monomicrobial supernatants. (C) *P. mirabilis* urease activity during incubation in an *E. faecalis* and *M. morganii* co-culture supernatants compared to a 50:50 mixture of monomicrobial supernatants. Dashed lines indicate OD_562_ of wells containing urine and urea in the absence of bacteria to control for pH change due to urea breakdown, error bars represent mean ± SD from a representative experiment with 3-6 replicates. (D) Urease activity is expressed as the mean change in optical density per minute (mOD/min) for the linear portion of the urease activity curves. The dashed lines indicates average urease activity for *P. mirabilis* during incubation in *P. mirabilis* Δ*ure* supernatant and the background level of pH change due to urea breakdown in the negative control wells. Error bars represent mean ± SD from three independent experiments with at least 3 replicates each. ^ns^*p*>0.05 and \*\*\**p*<0.001 by Student’s t-test.

*P. mirabilis* urease activity was significantly increased during incubation in cell-free spent urine supernatants from *E. faecalis, P. stuartii*, and *E. coli* compared to the level of activity for incubation in supernatant from the isogenic Δ*ure* mutant, and activity was significantly reduced during incubation in supernatant from *M. morganii* (Figure 2B and G). Therefore, modulation of *P. mirabilis* urease activity by each of these species is mediated by a soluble factor present in spent media and independent of direct cell-cell contact. To determine if either urease-modulatory activity is dominant, we next assessed urease activity in supernatants generated from co-cultures of each species with *M. morganii* compared to 50:50 mixtures of the monomicrobial supernatants. The urease activity curves for *E. faecalis* and *M. morganii* are shown in Figure 2C, and the other co-culture urease activity curves are shown in Fig S 1. Strikingly, all of the 50:50 mixtures resulted in an intermediate level of urease activity, while all of the co-culture supernatants dampened urease activity to a similar level as observed for the *M. morganii* monomicrobial supernatant (Figure 2C and G). Taken together, these results indicate that *M. morganii* is either able to prevent production of the enhancing signal by each of the other bacterial species, or that it inactivates, sequesters, or depletes the enhancing signal from the culture supernatant. In either case, *M. morganii* actively prevents enhancement of *P. mirabilis* urease activity by other bacterial species, in addition to directly dampening *P. mirabilis* urease activity.

### M. morganii prevents enhancement of P. mirabilis cytotoxicity by other bacterial species

Our prior studies with *P. mirabilis* and *P. stuartii* demonstrated that co-culture increased cytotoxicity to HEK293 cells *in vitro* compared to monomicrobial cultures, and that this process was not dependent upon urease activity from either species (47). We therefore utilized the *in vitro* cytotoxicity assay to determine if other bacterial species are capable of enhancing *P. mirabilis* cytotoxicity in a urease-independent manner, and if *M. morganii* can prevent enhancement. An example of the experimental approach is shown in Figure 3A. Briefly, each bacterial species of interest was incubated in filter-sterilized human urine for 90 minutes, washed twice, and overlaid onto a monlayer of HEK293 cells. After a 4-hour incubation, cytotoxicity was assessed by measurement of lactate dehydrogenase activity in the cell supernatants. For each polymicrobial condition, a predicted level of cytotoxicity (P) was calculated based on the proportion of the bacterial species in the culture and their relative cytotoxicity during monomicrobial culture. The level of cytotoxicity was then expressed relative to that of wild-type *P. mirabilis*, for ease of comparison between conditions.

**Figure 3.**
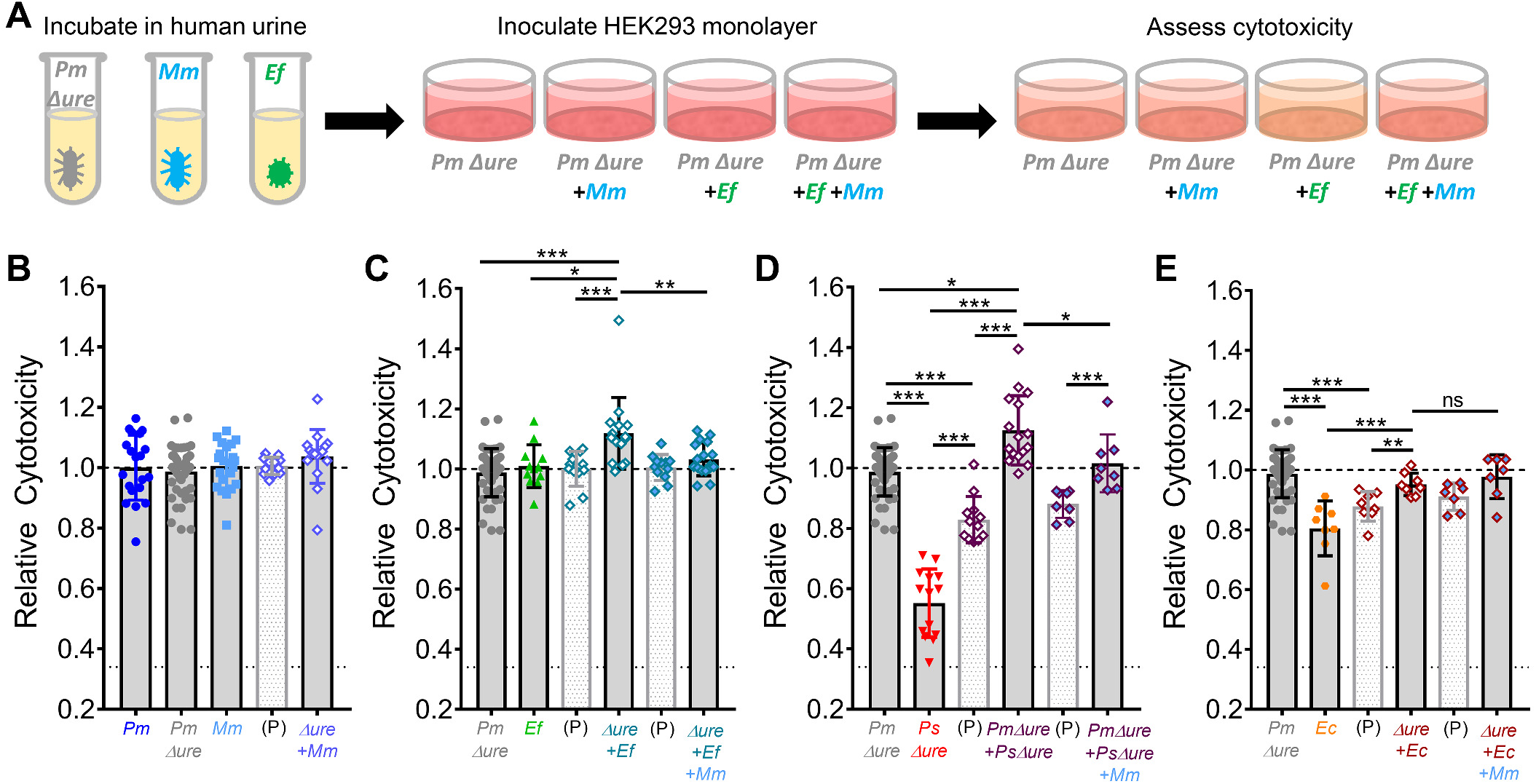
Urease-independent cytotoxicity of monomicrobial and polymicrobial cultures. Cytotoxicity to HEK293 cells was assessed by lactate dehydrogenase (LDH) release during a 4-hour incubation with various single-species, dual-species, and triple-species combinations as indicated. An example of the experimental approach is displayed in panel A. Bacteria were cultured in rich laboratory media to mid-log phase, centrifuged to pellet, resuspended in filter-sterilized human urine, and incubated for 90 minutes at 37°C with aeration prior to infection of HEK293 cells. The level of HEK293 lysis under each treatment condition is expressed relative to the level of treatment with *P. mirabilis* Δ*ure* (dashed line), and the dotted line indicates background cytotoxicity during treatment with sterile urine. For each dual-species and triple-species combination, a predicted (P) level of cytotoxicity was calculated based on the proportion of each bacterial species in the mixture and the cytotoxicity of their respective monomicrobial cultures. (B) Relative cytotoxicity of *P. mirabilis* HI4320, *P. mirabilis* Δ*ure, M. morganii*, and a dual-species culture of *P. mirabilis* Δ*ure* with *M. morganii.* (C) Relative cytotoxicity of *P. mirabilis* Δ*ure, E. faecalis*, a dual-species culture of *P. mirabilis* Δ*ure* with *E. faecalis*, and a triple-species culture of *P. mirabilis* Δ*ure* with *E. faecalis* and *M. morganii.* (D) Relative cytotoxicity of *P. mirabilis* Δ*ure, P. stuartii* Δ*ure*, a dual-species culture of *P. mirabilis* Δ*ure* with *P. stuartii* Δ*ure*, and a triple-species culture of *P. mirabilis* Δ*ure* with *P. stuartii* Δ*ure* and *M. morganii.* (E) Relative cytotoxicity of *P. mirabilis* Δ*ure, E. coli*, a dual-species culture of *P. mirabilis* Δ*ure* with *E. coli*, and a triple-species culture of *P. mirabilis* Δ*ure* with *E. coli* and *M. morganii.* Statistical significance was assessed by nonparametric Mann-Whitney test for indicated comparisons. ^ns^*p*>0.05, *\*p*<0.05, \*\**p*<0.01, \*\*\**p*<0.001.

Cytotoxicity was first compared for wild-type *P. mirabilis*, the isogenic urease mutant (*Pm* Δ*ure*), and *M. morganii* (Figure 3B). As expected, incubation with the urease mutant resulted in a similar level of cytotoxicity to HEK293 cells as wild-type *P. mirabilis*, confirming that urease does not contribute to cytotoxicity under these conditions. The urease mutant was therefore utilized for all further cytotoxicity experiments. Interestingly, *M. morganii* monomicrobial cultures exhibited a similar level of cytotoxicity as *P. mirabilis* Δ*ure*, and co-culture had no detectable impact on total cytotoxicity (Figure 3B).

*E. faecalis* monomicrobial cultures also exhibited comparable cytotoxicity as *P. mirabilis* Δ*ure*, but co-culture of *P. mirabilis* Δ*ure* with *E. faecalis* significantly increased cytotoxicity over the level observed for the monomicrobial cultures as well as the predicated value for the dual-species culture (Figure 3C). Thus, the interaction between these species appears to enhance production of other virulence factors in addition to increase urease activity. However, the addition of *M. morganii* dampened cytotoxicity back to a similar level as observed for each respective monomicrobial culture (Figure 3C). Taken together, these results indicate that co-culture of *P. mirabilis* with *E. faecalis* increases cytotoxicity in a urease-independent manner, and *M. morganii* dampens this enhancement.

Similarly, co-culture of *P. mirabilis* Δ*ure* with a *P. stuartii* urease mutant (*Ps* Δ*ure*) increased cytotoxicity above the level observed for either monomicrobial culture as well as the predicated co-culture level, which is consistent with our previous work (47). For this co-culture condition, *M. morganii* was capable of dampening enhancement back down to the level observed for *P. mirabilis* Δ*ure* alone, but not quite to the predicated level of activity (Figure 3D). In contrast, co-culture of *P. mirabilis* Δ*ure* with *E. coli* resulted in an intermediate level of cytotoxicity that was significantly greater than *E. coli* alone as well as the predicted value, but not greater than that of *P. mirabilis* Δ*ure* alone (Figure 3E). However, the addition of *M. morganii* had no discernable impact on cytotoxicity for this combination (Figure 3E). In summary, co-culture of *P. mirabilis* with *E. faecalis, P. stuartii*, or *E. coli* increases cytotoxicity to varying degrees through a urease-independent mechanism, and *M. morganii* is capable of preventing increased cytotoxicity from *E. faecalis* and *P. stuartii*, but not *E. coli*.

### M. morganii dampens disease severity during polymicrobial CAUTI

Our prior studies demonstrated that coinfection of *P. mirabilis* and *P. stuartii* increased disease severity compared to monomicrobial infection in a murine model of UTI and CAUTI. Specifically, enhancement of *P. mirabilis* urease activity by *P. stuartii* resulted in a greater incidence of urolithiasis and bacteremia (47, 48). We therefore hypothesized that coinfection of *P. mirabilis* with *E. faecalis* and *E. coli* may similarly increase disease severity, and that *M. morganii* may be capable of preventing progression to severe disease during polymicrobial infection.

To test this hypothesis, we utilized our robust animal model of CAUTI via the experimental approached detailed in Figure 4. Briefly, female CBA/J mice were transurethrally inoculated with 1×10^5^ CFUs of each species alone, or dual-species combinations as follows: i) *P. mirabilis* with *E. coli*, ii) *P. mirabilis* with *E. faecalis*, iii) *P. mirabilis* with *M. morganii*, and iv) *M. morganii* with *E. faecalis.* A triple-species infection was also conducted for *P. mirabilis* with *E. faecalis* and *M. morganii*, since co-cultures involving *E. faecalis* consistently presented the most severe phenotypes *in vitro.* For each mouse, a 4 mm segment of sterile silicone catheter tubing was inserted into the bladder at the time of inoculation to recapitulate CAUTI, as previously reported (47). Weight loss was monitored daily (Fig S 2), and mice were euthanized 96 hours post-inoculation (hpi) to quantify bacterial burden (Figure 5 and Fig S 3) and the incidence of bacteremia and macroscopic kidney stones (Table 1). For each infection group, a subset of mice had the entire bladder and both kidneys sectioned and paraffin embedded for assessment of histopathology rather than quantification of bacterial burden (Figure 5F-H and Fig S 4).

**Figure 4.**
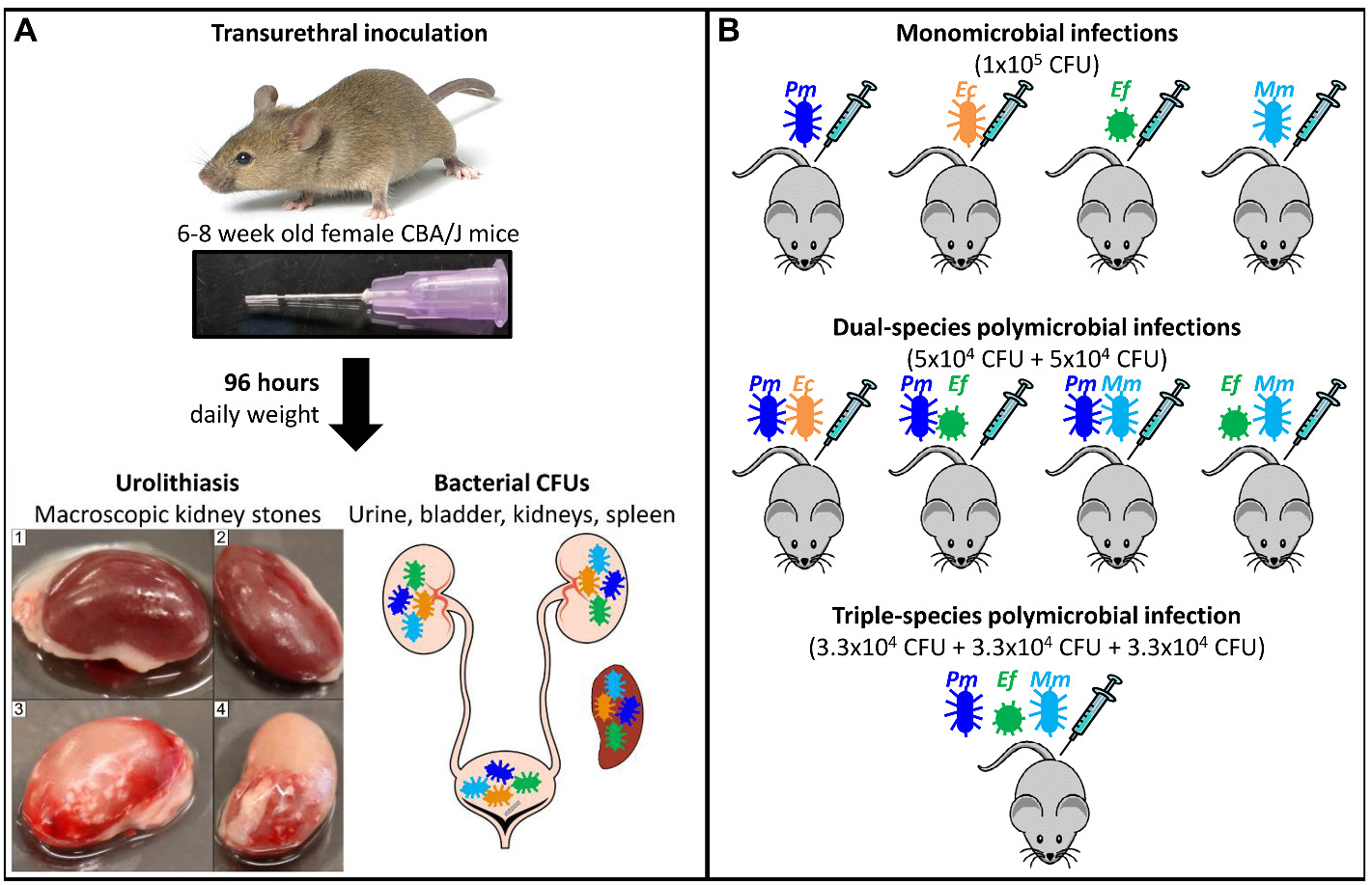
Experimental schematic for murine CAUTI. (A) Overview of experimental approach for murine CAUTI. Female CBA/J mice were purchased from Jackson laboratories at 5-7 weeks of age, allowed to acclimate to the vivarium for 1 week, and infected at 6-8 weeks of age by transurethral instillation of 50µl of a bacterial inoculum over 25-seconds. At the time of inoculation, a 4mm segment of sterile silicone catheter tubing was inserted into the bladder. All mice were weighed daily in the morning to monitor weight loss. After 96 hours, urine was collected from all mice for determination of bacterial burden, mice were euthanized, and urolithiasis was assessed by the presence of macroscopic kidney stones. Images 1 and 2 display infected kidneys lacking stones, while images 3 and 4 demonstrate the presence of severe macroscopic kidney stones. Bladders, kidneys, and spleens were also removed for determination of bacterial burden. (B) Overview of bacterial inoculations to establish monomicrobial and polymicrobial infections. All mice were inoculated with 1×10^5^ colony forming units (CFU) of bacteria. *Proteus mirabilis* HI4320 (*Pm*, blue), *Escherichia coli* CFT073 (*Ec*, orange), *Enterococcus faecalis* 3143 (*Ef*, green), and *Morganella morganii* (*Mm*, light blue).

**Figure 5.**
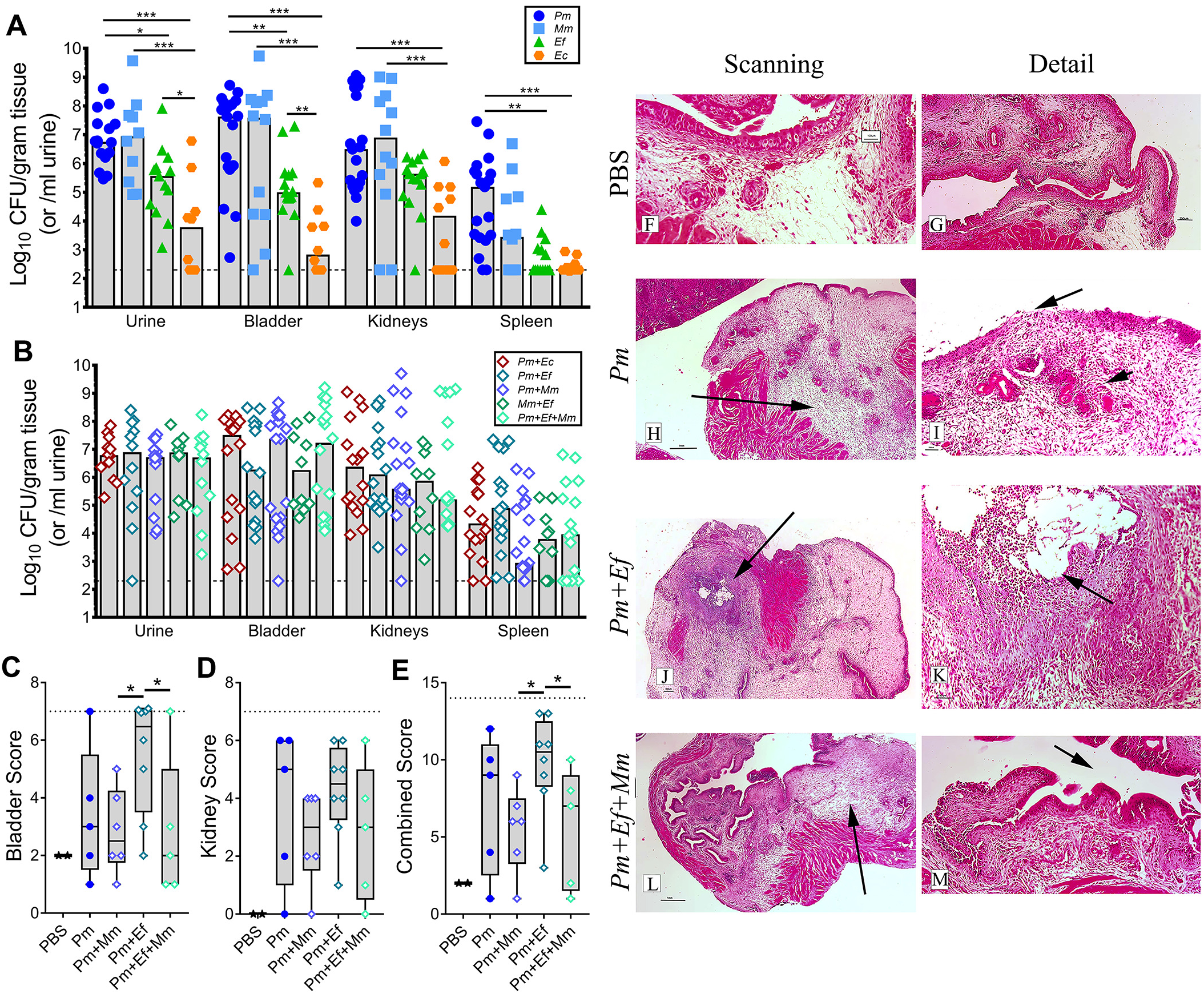
Urinary tract bacterial burden and tissue histopathology during monomicrobial and polymicrobial CAUTI. The log_10_ CFUs recovered from the urine, bladder, kidneys, and spleen per gram of tissue or milliliter of urine are shown for monomicrobial infections (A), as well as the total CFUs recovered from polymicrobial infections (B). Each symbol represents the CFUs recovered from a single mouse, gray bars indicate the median, and dashed lines indicate the limit of detection (200 CFU). The number of mice in each infection group is provided in Table 1. For statistical analysis, all CFU data were log_10_-transformed and significance was assessed by 2-way ANOVA with Tukey’s multiple comparisons test *\*p*<0.05, \*\**p*<0.01, \*\*\**p*<0.001. (C-M) Bladder and kidney sections from mice inoculated with PBS (2), *P. mirabilis* (5), *P. mirabilis* + *M. morganii* (6), *P. mirabilis* + *E. faecalis* (8), or *P. mirabilis* + *E. faecalis* + *M. morganii* (5) were paraffin embedded, sectioned, and stained with hematoxylin and eosin for assessment of tissue damage and inflammation using the scoring rubric provided in Supplemental Table 1. (C) Bladder scores, (D) kidney scores, (E) combined score for the bladder and kidney from each individual mouse. *\*p*<0.05 by nonparametric Mann-Whitney test. (F-M) Representative bladder images from select infection groups, including a low-magnification (2x objective) scanning view (F, H, J, L) and a higher magnification detail view (G, I, K, M). (F and G) PBS/mock infected mice, normal healthy bladder. (H) *P. mirabilis* monomicrobial infection with severe edema (arrow) and multifocal vasculitis. (I) Detail of vasculitis (arrowhead), widespread neutrophilic inflammation, and epithelial ulcer (arrow). (J) *P. mirabilis + E. faecalis* infection showing severe edema, neutrophilic inflammation, and ulceration with embedded calculus (arrow). (K) Detail of calculus (arrow) and surrounding inflammation and necrosis. (L) *P. mirabilis + E. faecalis + M. morganii* infection showing intact mucosa and moderate edema (arrow). (M) Detail of intact mucosa (arrow).

**Table 1.** CAUTI disease severity.

| **Table 1** | **Number of mice** | **Bacteremia<sup>A</sup>** | ***P* value** | **Urolithiasis<sup>B</sup>** | ***P* value** |
| --- | --- | --- | --- | --- | --- |
| *Monomicrobial Infections* |  |  |  |  |  |
| *Ec* | 10 | 20% | .001* | 0% | --- |
| *Ef* | 15 | 47% | .040* | 0% | --- |
| *Mm* | 15 | 62% | .222 | 19% | .666 |
| *Pm* | 27 | 78% | --- | 26% | --- |
| *Polymicrobial Infections* |  |  |  |  |  |
| *Pm+Ec* | 17 | 82% | .714 | 29% | .800 |
| *Pm+Ef* | 25 | 96% | .054 | 56% | .027* |
| *Pm+Ef+Mm* | 21 | 71% | .614 | 24% | .867 |
| *Pm+Mm* | 26 | 73% | .691 | 15% | .344 |
| *Ef+Mm* | 10 | 70% | .624 | 10% | .296 |
| *Urease-Independent Infections* |  |  |  |  |  |
| *Pm Δure* | 10 | 40% | .001* | 0% | --- |
| *Δure+Ec* | 10 | 40% | .999 | 0% | --- |
| *Δure+Ef* | 12 | 58% | .392 | 0% | --- |
| *Δure+Mm* | 8 | 37% | .914 | 12% | --- |
| *Δure+Ef+Mm* | 12 | 33% | .746 | 8% | --- |

<sup>A</sup>Bacteremia was assessed by spleen CFUs above the limit of detection. <sup>B</sup>Urolithiasis was assessed by the presence of macroscopic kidney stones, as shown in Figure 3. Severity is expressed relative to *P. mirabilis* (*Pm*, white), with more severe being orange and less severe being blue. *P* values determined by chi-square *versus* to *Pm* monomicrobial infection for all groups except the Urease-Independent Infections, which were compared to *Pm Δure*.

All mice exhibited weight loss during the first 24 hpi, regardless of infection group, and the majority of mice failed to recover weight during the study. The only group of mice that exhibited recovery from the initial weight loss were those that received an *E. coli* monomicrobial inoculum (Fig S 2A, C, and H). This is likely because monomicrobial inoculation with 1×10^5^ CFUs of *E. coli* resulted in the lowest bacterial burden of all of the infection groups (Figure 5A) as well as the lowest overall infection severity (Table 1), followed by the *E. faecalis* monomicrobial infection. No other significant differences in weight loss were observed.

Despite differences in the colonization density achieved by each species during monomicrobial infection (Figure 5A), there were no significant differences in total bacterial burden within the urinary tract during any of the polymicrobial infections (Figure 5B). The CFUs recovered for each individual bacterial species from the polymicrobial infections are shown across infection groups in Fig S 4A-D and on a per-mouse basis Fig S 4E-I. *P. mirabilis* CFUs and *E. faecalis* CFUs were stable across all infection types (Fig S 4A and B), *M. morganii* CFUs were reduced in the presence of *P. mirabilis* but unaffected by *E. faecalis* (Fig S 4C), and *E. coli* CFUs were augmented by *P. mirabilis* (Fig S 4D). Despite these shifts in bacterial burden for each species, the total combined CFUs achieved within the urinary tract were consistent across all of the polymicrobial infections (Figure 5B).

The similarities in total bacterial burden across infection groups is particularly noteworthy considering that there were dramatic differences in disease severity between infection groups (Table 1). *P. mirabilis* exhibited the greatest disease severity of the monomicrobial infections, with 78% of mice exhibiting bacteremia and 26% urolithiasis. Coinfection of *P. mirabilis* with *E. coli* did not significantly impact the incidence of bacteremia or urolithiasis compared to *P. mirabilis* monomicrobial infection, despite the ability of this combination to increase urease activity and cytotoxicity *in vitro*. In contract, coinfection of *P. mirabilis* with *E. faecalis* resulted in the most severe disease presentation, as 96% of these mice exhibited bacteremia (+18% relative to *P. mirabilis* monomicrobial infection, *p*=0.054) and 56% exhibited urolithiasis (+30% relative to *P. mirabilis* monomicrobial infection, *p*=0.027). This coinfection combination also resulted in the greatest incidence of maximum histopathology scores for tissue damage and inflammation in the bladder (Figure 5C) and the highest combined scores for bladder and kidney damage (Figure 5E). Representative bladder sections are displayed in Figure 5F-M, and representative kidney sections are displayed in Fig S 4.

Remarkably, the addition of *M. morganii* to the *P. mirabilis* and *E. faecalis* coinfection dramatically reduced infection severity to level that was indistinguishable from mice infected with *P. mirabilis* alone or mice coinfected with *P. mirabilis* and *M. morganii* (Table 1). Seventy-one percent of mice from the triple-species infection exhibited bacteremia (−25% compared to *P. mirabilis* + *E. faecalis* coinfection, *p*=0.0208) and 24% exhibited urolithiasis (−32% compared to *P. mirabilis* + *E. faecalis* coinfection, *p*=0.0272). The decrease in disease severity was also clearly evident by histopathology, as the triple-species infection resulted in significantly reduced bladder scores and combined scores than the *P. mirabilis* + *E. faecalis* coinfection (Figure 5C and E). Thus, the ability of *M. morganii* to dampen enhancement of urease activity and cytotoxicity *in vitro* does indeed translate to a reduction in infection severity *in vivo*. Taken together, these data indicate that disease severity during polymicrobial CAUTI is dependent on the effects of the specific co-infecting organisms, rather than bacterial burden alone, and that an organism such as *M. morganii* can reduce infection severity even when present at a low density.

### Modulation of polymicrobial infection severity is predominantly mediated through a urease-dependent mechanism

Our prior studies of *P. mirabilis* with *P. stuartii* demonstrated that increased disease severity during coinfection of these species was dependent on the presence of a functional urease operon in *P. mirabilis* and enhancement of urease activity (47, 48). We therefore tested the hypothesis that *P. mirabilis* urease activity is required for modulation of disease severity during polymicrobial infection. Mice were inoculated as above, only using the isogenic urease mutant of *P. mirabilis* HI4320, Δ*ure*. Fig S 5 shows percent weight change. Consistent with the prior infections, all mice exhibited weight loss during the first 24 hpi, regardless of infection group. However, as expected, the majority of mice inoculated with Δ*ure* exhibited decreased disease severity compared to infection with the wild type strain, as indicated by a trend towards weight gain from 72-96 hpi.

Figure 6 shows the total CFUs recovered from mice inoculated with *P. mirabilis* Δ*ure* alone for comparison to each of the polymicrobial infections. The CFUs recovered for each individual bacterial species from the polymicrobial infections are shown across infection groups in Fig S 6A-D and on a per-mouse basis Fig S 6E-H. Similar to infections with wild-type *P. mirabilis*, there were no significant differences in total bacterial burden within the urinary tract between infection groups (Figure 6). *P. mirabilis* Δ*ure* CFUs were again largely similar across all infection types (Fig S 6A). Interestingly, *E. faecalis* CFUs were significantly decreased in the urine and kidneys when *P. mirabilis* Δ*ure* was present (Fig S 6B), which was not observed during infection with the wild-type strain (Fig S 3D), suggesting that *E. faecalis* colonization may be perturbed by *P. mirabilis* in the absence of urease activity. *M. morganii* colonization was again significantly decreased in the urine, bladder, and kidneys when *P. mirabilis* Δ*ure* was present, indicating that the impact of *P. mirabilis* on *M. morganii* colonization is urease-independent (Fig S 6C). Similarly, *E. coli* colonization was significantly increased when *P. mirabilis* Δ*ure* was present, indicating that urease activity is not required for enhancement of *E. coli* colonization during coinfection (Fig S 6D).

**Figure 6.**
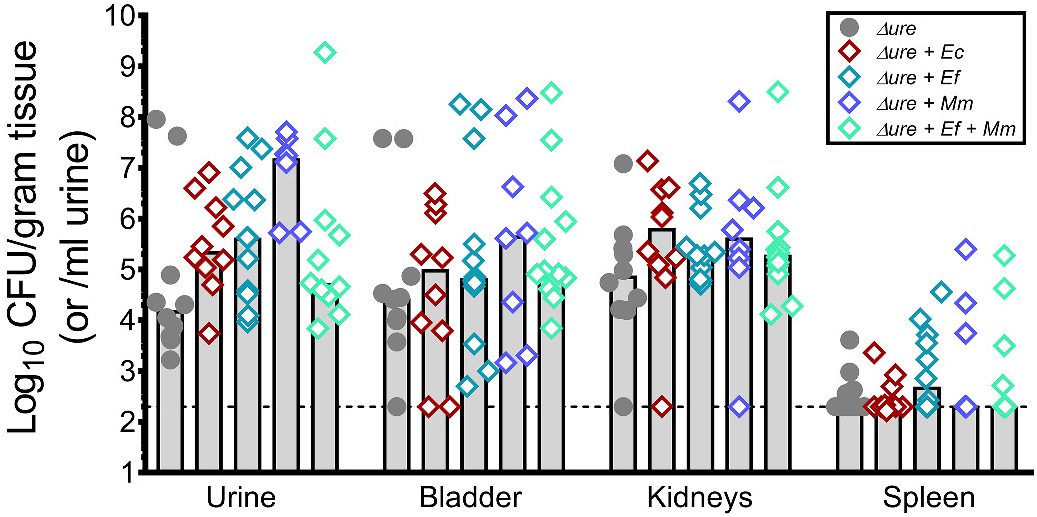
Urinary tract bacterial burden during monomicrobial and polymicrobial infection with *P. mirabilis* Δ*ure*. The log_10_ CFUs recovered from the urine, bladder, kidneys, and spleen per gram of tissue or milliliter of urine are displayed for *P. mirabilis* Δ*ure* monomicrobial infection as well as the total CFUs recovered from polymicrobial infections. Each symbol represents the CFUs recovered from a single mouse, gray bars indicate the median, and dashed lines indicate the limit of detection (200 CFU). The number of mice in each infection group is provided in Table 1. All CFU data were log_10_-transformed, and no significant differences were identified by 2-way ANOVA with Tukey’s multiple comparisons test.

Despite these differences in the colonization density achieved by each individual bacterial species during co-infection, the total bacterial burden with the urinary tract again remained consistent across infection groups (Figure 6). Compared to monomicrobial infection with wild-type *P. mirabilis*, Δ*ure* infections exhibited a significant reduction in the incidence of bacteremia (40% compared to 78%, *p*<0.001), which is consistent with the role of urease activity and urolithiasis in tissue damage and progression to bacteremia (47, 48). Even more striking was the observation that disease severity remained consistent between monomicrobial infection with *P. mirabilis* Δ*ure* and each of the polymicrobial infections, in stark contrast to the results obtained with wild-type *P. mirabilis* (Table 1). Thus, while urease-independent interactions still have the potential to influence disease severity during polymicrobial *P. mirabilis* CAUTI, modulation of urease activity is the main contributor to infection severity in this model system. The mechanism(s) by which *M. morganii* dampens *P. mirabilis* urease activity and prevents enhancement by other bacterial species may therefore elucidate potent inhibitors of infection severity, and could have clinical utility for CAUTIs involving *P. mirabilis*.

## Discussion

It is now widely recognized that microbes generally exist in complex polymicrobial communities rather than living in isolation. There are numerous experimental and clinical examples of polymicrobial interactions increasing disease severity compared to monomicrobial infection (17–20), as well as examples of interactions that attenuate disease severity (21). However, the clinical significance of polymicrobial interactions are less clear in the context of bacteriuria, particularly for individuals with indwelling urinary catheters. In the present study, we show that bacterial species colonizing the catheterized urinary tract can both enhance (*E. faecalis, E. coli*, and *P. stuartii*) and dampen (*M. morganii*) disease severity by exerting an effect on *P. mirabilis*, and that this process does not require physical contact or close proximity of the bacteria. We further demonstrate that the predominant cause of disease severity is modulation of *P. mirabilis* urease activity, but additional urease-independent factors also contribute to cytotoxicity to host cells.

*M. morganii* is a common constituent of the intestinal tract and considered to be a rare opportunistic pathogen, mainly causing nosocomial urinary tract infections in elderly catheterized individuals (53). This species has occasionally been identified as a cause of severe UTI-related complications such as sepsis and bacteremia, although it is less common than *P. mirabilis* or *E. coli* and more likely to be present during polymicrobial sepsis than as the sole causative agent of disease (54). Thus, severe disease manifestations such as entry of *M. morganii* to the bloodstream is likely facilitated by other bacterial species, particularly within the urinary tract where it is generally present during polymicrobial infection (6, 55). Patients experiencing *M. morganii* sepsis are also more likely to have multiple comorbidities and reduced functional status compared to those with *E. coli* (54), underscoring the opportunistic nature of this organism. It is therefore fascinating that this rare opportunist has such a dramatic impact on the pathogenesis of polymicrobial CAUTI involving *P. mirabilis.* Our results clearly demonstrate that *M. morganii* dampens *P. mirabilis* urease activity and even prevents enhancement by other bacterial species, which is consistent with prior reports of *M. morganii* reducing catheter blockage and encrustation by *P. mirabilis in vitro* (24). However, our study further extends these observations to show that *M. morganii* significantly attenuates polymicrobial CAUTI.

Our experimental findings have important clinical implications, particularly for patient populations requiring long-term catheterization. Approximately 50% of individuals catheterized for 28 days or longer experience catheter blockage and/or encrustation from crystalline deposits, the vast majority of which are the result of *P. mirabilis* urease activity (10, 12, 56, 57). These crystalline deposits accumulate on the catheter surface and facilitate biofilm formation (58), a bacterial mode of growth that is associated with immune evasion and antimicrobial resistance (59). Many of the urease-positive organisms that colonization urinary catheters can persist within a catheterized host for months, even after antibiotic treatment and catheter changes (14). Thus, interactions between bacteria within the catheterized bladder and in catheter biofilms have the potential to significantly impact catheter encrustation, tissue damage, urolithiasis, and dissemination to the kidneys and bloodstream.

The implications of our findings, in this context, are depicted in Figure 7. When *P. mirabilis* is present in the bladder in the absence of other bacterial species, it is capable of causing stone formation, crystalline catheter biofilms, ascending to the kidneys, and potentially dissemination to the bloodstream (Figure 7A). When an enhancing species is present, such as *P. stuartii, E. coli*, or *E. faecalis*, secreted factors produced by these organisms act on *P. mirabilis* to increase urease activity and cytotoxicity. This leads to an increase in tissue damage, urine pH and crystalline deposits, and catheter encrustation and blockage, in addition to promoting urolithiasis and bacteremia. Bladder co-colonization of *P. mirabilis* with *E. coli* also bolsters *E. coli* CFUs, which may potentiate its impact on *P. mirabilis* (Figure 7B). However, when *M. morganii* is present, it secretes factors that counteract the effects of these other microbes on *P. mirabilis* urease activity and cytotoxicity, thereby reducing crystal formation, catheter encrustation and blockage, urolithiasis, and bacteremia (Figure 7C). Considering the level of activity present in cell-free urine supernatants of these organisms, it is likely that if any of these modulatory organisms are present in the bladder, they would exert an impact on catheter-resident *P. mirabilis*.

**Figure 7.**
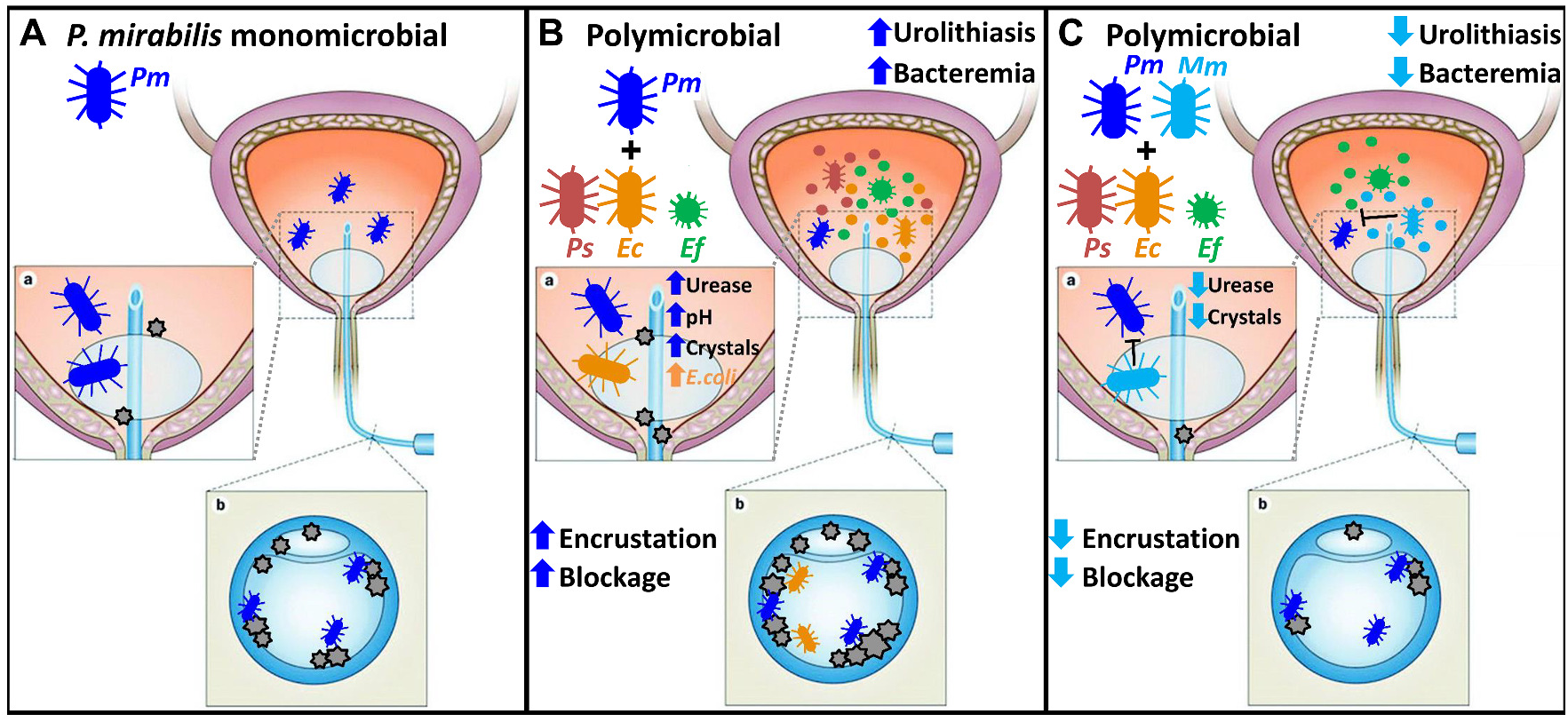
Implications of polymicrobial interactions during CAUTI. (A) *P. mirabilis* (*Pm*, blue) monomicrobial colonization. Urease activity from this bacterial species results in crystalline deposits in the bladder (a, bladder inset) and on the catheter (b, catheter inset), promoting bacterial colonization and in some cases bacteremia. (B) Polymicrobial colonization involving *P. mirabilis* and either *P. stuartii* (*Ps*, maroon), *E. coli* (*Ec*, orange), or *E. faecalis* (*Ef*, green). Co-colonizing species secrete factors that act on *P. mirabilis* to enhance urease activity, thereby raising urine pH and increasing crystal formation, which increases the risk of catheter encrustation and blockage. Co-colonization with *E. faecalis* also increases the risk of urolithiasis and bacteremia, and *E. coli* colonization is augmented by the presence of *P. mirabilis*. (C) Polymicrobial colonization involving *P. mirabilis* and *M. morganii* (*Mm*, light blue) with *P. stuartii, E. coli*, or *E. faecalis*. *M. morganii* secretes factors that act on *P. mirabilis* to dampen urease activity and prevent enhancement by other species, such as *E. faecalis*, thereby reducing the formation of crystals and the risk of catheter encrustation and blockage. The presence of *M. morganii* also decreases risk of urolithiasis and bacteremia during coinfection with *P. mirabilis* and *E. faecalis*. Catheterized bladder image adapted from (78, 79).

A handful of other dual-species polymicrobial combinations have been characterized in murine models of UTI or CAUTI, although most studies have focused on the impact of coinfection on colonization rather than disease severity. For instance, *E. faecalis* and *Streptococcus agalactiae* are both capable of promoting *E. coli* colonization by suppressing the host immune response (60, 61). Coinfection of *E. coli* with *P. mirabilis* promotes colonization of both species during uncomplicated UTI through a dampening of the immune response (62), although only *E. coli* received a benefit from co-colonization in our CAUTI model. Coinfection of *P. mirabilis* with *P. stuartii* has no impact on bacterial burden of either species within the urinary tract, but their interactions actually promotes a potent proinflammatory response, greater tissue damage, and an increased incidence of severe disease during both UTI and CAUTI (47, 48). This is likely also the case for coinfection of *P. mirabilis* with *E. faecalis*, given the increase in cytotoxicity that occurs *in vitro* and the incidence of bacteremia in our CAUTI model. Increased likelihood of severe disease has also been observed during coinfection of *P. mirabilis* with *Staphylococcus saprophyticus* (63), and for *P. aeruginosa* with *E. faecalis* (64).

In contrast to these infection-promoting interactions, there are limited examples of polymicrobial interactions that attenuate disease severity during UTI or CAUTI. Production of an iron-binding factor by commensal *E. coli* and other *Enterobacteriaceae* antagonizes iron acquisition by *P. aeruginosa*, thereby limiting growth and possibly survival in the catheterized bladder (65). Other commensal *E. coli* strains have also been shown to inhibit growth and catheter colonization by traditional uropathogens *in vitro* (66, 67). Thus, our observation of disease modulation by *M. morganii* is consistent with the idea that commensal or non-traditional pathogens can hinder establishment of infection and disease severity by other pathogens.

In conclusion, our results indicate that further assessments of the clinical significance of polymicrobial bacteriuria and CAUTI need to consider the exact combination of bacterial species that are present, in addition to Gram-positive *vs.* Gram-negative and bacterial burden within the sample. Such interactions may be difficult to capture from single urine samples if colonization of the disease-modulating organism is impacted during coinfection, as is the case for *M. morganii* and *E. coli.* However, a deeper understanding of the importance of these interactions is necessary to help guide novel treatment strategies. Furthermore, identification of *M. morganii* secreted products that modulate *P. mirabilis* pathogenicity has the potential to lead to new therapeutic treatments for CAUTI. The abundance of urease-dampening and urease-enhancing factors during asymptomatic colonization may also have predictive value for patient outcomes in long-term catheterization.

## Materials and Methods

### Study Approval

Animal protocols were approved by the Institutional Animal Care and Use Committee (IACUC) of the University at Buffalo (MIC31107Y), in accordance with the Office of Laboratory Animal Welfare (OLAW), the United States Department of Agriculture (USDA), and the Association for Assessment and Accreditation of Laboratory Animal Care (AAALAC). Mice were anesthetized with a weight-appropriate dose (0.1 ml for a mouse weighing 20 gm) of ketamine/xylazine (80-120 mg/kg ketamine and 5-10 mg/kg xylazine) by IP injection, and euthanized by inhalant anesthetic overdose with vital organ removal.

### Bacterial strains and growth conditions

*Proteus mirabilis* HI4320, *Providencia stuartii* BE2467, and *Morganella morganii* TA43 were isolated from the urine of catheterized patients in a chronic care facility in Maryland (12). *Escherichia coli* CFT073 was isolated from a patient hospitalized for acute pyelonephritis (68). *Enterococcus faecalis* 3143 was isolated from the urine of a catheterized nursing home resident in Michigan (47). *P. mirabilis* Δ*ure* refers to *P. mirabilis* HI4320 lacking urease activity due to an insertional-disruption of the *ureF* gene, and *P. stuartii* Δ*ure* refers to *P. stuartii* BE2467 lacking the portion of its plasmid that encodes the urease operon (47, 69, 70). Bacteria were routinely cultured at 37°C with aeration in 5 ml LB broth (10 g/L tryptone, 5 g/L yeast extract, 0.5 g/L NaCl) or on LB solidified with 1.5% agar. *Enterococcus faecalis* isolates were cultured in brain heart infusion (Difco) broth. LB was supplemented with 25 µg/ml kanamycin, 20 µg/ml chloramphenicol, 25 µg/ml ampicillin, 2.5 µg/ml tetracycline, or 100 µg/ml streptomycin to distinguish between species.

### Urine growth curves

Pooled, filter-sterilized human urine from female donors (Cone Bioproducts) was diluted 1:1 with sterile saline to achieve a uniform specific gravity of ∼1.005. Overnight cultures of bacterial species were diluted 1:100 into 5 ml of urine for monomicrobial growth curves. For co-cultures, each bacterial species was diluted 1:200 to maintain the same total starting CFUs, and for triple cultures each bacterial species was diluted 1:300. Bacterial burden was assessed by plating for CFUs. To distinguish between bacterial species, samples were plated as follows: 1) *P. mirabilis + E. faecalis* coinfection samples were plated on plain LB (total CFUs) and BHI with streptomycin (*E. faecalis* CFUs); 2) *P. mirabilis + M. morganii* coinfection samples were plated on plain LB (total CFUs) and LB with ampicillin (*M. morganii* CFUs), 3) *P. mirabilis* + *E. coli* coinfection samples were plated on plain LB (total CFUs) and LB with tetracycline (*P. mirabilis* CFUs), 4) *E. faecalis* + *M. morganii* coinfection samples were plated on plain LB (total CFUs) and BHI with streptomycin (*E. faecalis*), and 5) *P. mirabilis* + *E. faecalis* + *M. morganii* triple infection samples were plated on plain LB (total CFUs), LB with ampicillin (*M. morganii* CFUs), and BHI with streptomycin (*E. faecalis* CFUs).

### Urease assay

A previously described alkalimetric screen was used to measure urease activity in whole bacterial cells (47). Briefly, bacterial strains were cultured to mid-log phase (OD_600_=0.5), centrifuged to pellet, and resuspended in filter-sterilized human urine (using a batch pool from de-identified donors, purchased from Cone Bioproducts). Resuspended cultures were incubated individually at 37°C with aeration or mixed in equal proportions to generate co-cultures. After 90 minutes, cultures were centrifuged to pellet, and supernatants were passaged through a 0.22µm pore size filter to generate cell-free spent urine supernatants for determination of *P. mirabilis* urease activity. For measurement of urease enhancement in spent urine supernatants, *P. mirabilis* HI4320 was cultured for ∼16 hours 37°C with aeration in 5 ml LB broth, centrifuged to pellet, and resuspended in 0.9% sterile saline. The urine supernatants were dispensed into replicated wells of a 96-well plate, supplemented with a phenol red and urea mixture to a final concentration of 0.001% WT/vol phenol red and 500 mM urea, and inoculated with 20 μl of the bacterial saline suspension. Optical density at 562 nm was measured every 60 seconds for 2 hours using a Synergy H1 (BioTek). Urease activity was expressed as the mean change in optical density per minute (mOD/min) for the linear portion of the activity curve, as calculated by the Gen5 software (BioTek).

### Cytotoxicity

Toxicity of bacterial species independently or during co-culture were estimated by the lactate dehydrogenase (LDH) release cytotoxicity assay as previously described (47, 71). HEK293 cells were purchased from ATCC in 2018 and used for cytotoxicity studies after ≤ 3 passages. Briefly, bacteria were cultured in LB to mid-log phase (OD_600_=0.5), centrifuged to pellet, and resuspended in filter-sterilized human urine to induce urease activity. Resuspended cultures were incubated for 60 minutes at 37°C. Cultures were then pelleted, washed twice, resuspended in DMEM without supplements or antibiotics to a concentration of 2×10^7^ CFU/ml, and added to a monolayer of low-passage HEK293 cells at a multiplicity of infection of approximately 100:1. After a 4 hour incubation at 37°C and 5% CO_2_, the amount of LDH released into the supernatant was measured using an LDH Cytotoxicity assay kit (Cayman Chemical). Treatment with 9% Triton X-100 for the 4 hour incubation was used as a positive control for maximum lysis, and treatment with sterile urine alone was used as a negative control. The level of HEK293 lysis under each treatment condition is expressed relative to the level for *P. mirabilis* Δ*ure*. For all dual-species and triple-species cultures, a predicted (P) level of cytotoxicity was calculated based on the proportion of each bacterial species in the culture (determined by differential plating for CFUs) and the relative cytotoxicity of monomicrobial cultures for each species.

### Mouse model of CAUTI

Infection studies were carried out as previously described (47). Briefly, bacteria were cultured overnight in LB, washed in phosphate-buffered saline (PBS: 0.128 M NaCl, 0.0027 M KCl, pH 7.4), adjusted to 2×10^8^ CFU/ml (OD_600_=0.2 for *P. mirabilis, P. stuartii*, and *M. morganii*, and OD_600_=0.4 for *E. coli* and *E. faecalis*), and diluted 1:100 to achieve an inoculum of 2×10^6^ CFU/ml. Female CBA/J mice at 6-8 weeks of age (Jackson) were inoculated transurethrally with 50 µl of 2×10^6^ CFU/ml (1×10^5^ CFU/mouse) of either a single bacterial species (monomicrobial), a 1:1 mixture of two species for coinfections, or a 1:1:1 mixture of *P. mirabilis* with two other species for polymicrobial infections. In all cases, a 4 mm segment of sterile silicone tubing (0.64 mm O.D., 0.30 mm I.D., Braintree Scientific, Inc.) was carefully advanced into the bladder during inoculation as described elsewhere (47, 72–74). Mice were euthanized 96 hours post-inoculation (hpi) and bladders, kidneys and spleens were harvested into 5 mL Eppendorf tubes containing 1 mL PBS. Gross inspection of kidneys was performed to assess macroscopic urolithiasis. Tissues were homogenized using a Bullet Blender 5 Gold (Next Advance) and plated using an EddyJet 2 spiral plater (Neutec Group) for determination of CFUs using a ProtoCOL 3 automated colony counter (Synbiosis). To distinguish between bacterial species, samples were plated as follows: 1) *P. mirabilis + E. faecalis* coinfection samples were plated on plain LB (total CFUs) and BHI with streptomycin (*E. faecalis* CFUs); 2) *P. mirabilis + M. morganii* coinfection samples were plated on plain LB (total CFUs) and LB with ampicillin (*M. morganii* CFUs), 3) *P. mirabilis* + *E. coli* coinfection samples were plated on plain LB (total CFUs) and LB with tetracycline (*P. mirabilis* CFUs), 4) *E. faecalis* + *M. morganii* coinfection samples were plated on plain LB (total CFUs) and BHI with streptomycin (*E. faecalis*), and 5) *P. mirabilis* + *E. faecalis* + *M. morganii* triple infection samples were plated on plain LB (total CFUs), LB with ampicillin (*M. morganii* CFUs), and BHI with streptomycin (*E. faecalis* CFUs).

### Pathological evaluation

A subset of mice from each infection group were selected for histopathological examination rather than determination of bacterial burden. For these mice, bladders were cut longitudinally and each kidney was transversely. Organs were preserved in 10% formalin, embedded in paraffin, sectioned, and stained with hematoxylin and eosin. Sections were examined microscopically and scored in a blinded fashion by a veterinary pathologist to determine the severity and extent of inflammation and lesions using an adaptation of a previously developed semi-quantitative scoring system (47, 71, 75–77). The scoring rubric is provided in Table S 1.

### Statistics

Normalcy was assessed for all datasets by the Shapiro-Wilk and D’Agostino & Pearson normality tests. Significance was assessed using two-way analysis of variance (ANOVA), nonparametric Mann-Whitney test, unpaired *t* test, or chi square test, as indicated in the figure legends. All *P* values are two tailed at a 95% confidence interval. All analyses were performed using GraphPad Prism, version 7.03 (GraphPad Software, San Diego, CA).

## Acknowledgments

We would like to thank members of the Department of Microbiology & Immunology in the Jacobs School of Medicine and Biomedical Sciences at the University at Buffalo for helpful comments and critiques. This work was supported by the National Institutes of Health [R00 DK105205 to C.E.A.]. The sponsors were not involved in the study design, methods, subject recruitment, data collections, analysis, or preparation of the paper. The content is solely the responsibility of the authors and does not necessarily represent the official views of the funders.

